# Pervasive co-option of prokaryotic adenine methyltransferases by eukaryotic retrotransposons

**DOI:** 10.64898/2026.05.21.726589

**Authors:** Manuel Fernández-Moreno, Pedro Romero Charria, Lan Xu, Beatriz Loria-Vinueza, Manuel Irimia, Ignacio Maeso, Alex de Mendoza

## Abstract

Cytosine DNA methylation is broadly associated with transposable element silencing across eukaryotes, whereas 6-methyladenine (6mA) in unicellular eukaryotes is linked to actively transcribed chromatin. How transposable elements adapt to these contrasting epigenetic environments remains largely unexplored. Here we identify widely distributed eukaryotic retrotransposons encoding prokaryotic-like DNA adenine methyltransferases (DAMs). Phylogenetic analyses indicate a single ancestral acquisition from prokaryotes followed by recurrent transfers between retrotransposon classes across diverse eukaryotes. DAM-carrying LTR elements are preferentially found in species encoding AMT1, the main eukaryotic 6mA methyltransferase, and show elevated 6mA levels relative to other LTR retrotransposons in multiple lineages, accompanied by increased transcription. We further identify retrotransposons combining adenine and cytosine methyltransferases with chromodomains, indicating the assembly of unexpectedly complex epigenetic toolkits within single retrotransposon units. These findings suggest that retrotransposons have repeatedly co-opted prokaryotic-like methyltransferases to exploit host 6mA-associated chromatin, highlighting adaptation to host epigenetic landscapes as a major driver of transposable element evolution.

## INTRODUCTION

5-methylcytosine (5mC) is a widespread DNA modification in eukaryotes and plays a major role in the silencing of transposable elements (TEs) across diverse lineages ^1,2^. In contrast, the presence and function of N6-methyladenine (6mA) in eukaryotes has remained more controversial. Although several reports of 6mA in multicellular organisms have been challenged^3–5^, an emerging consensus supports 6mA as an ancestral eukaryotic DNA modification that is retained in many unicellular eukaryotes but lost from all major multicellular lineages^6^. In the unicellular species that retain it, 6mA is typically enriched at transcription start sites of actively transcribed genes and co-occurs with permissive chromatin states, including H3K4me3-marked nucleosomes^6–8^. This pattern is strongly associated with the presence of the adenine DNA methyltransferase AMT1, which maintains symmetrical ApT methylation^7,9^. Unlike 5mC, which is commonly linked to TE repression, 6mA is generally depleted from transposable elements^6^. However, whether this exclusion applies broadly across repetitive elements, or whether some TEs have evolved mechanisms to exploit 6mA to promote their own activity, remains completely unexplored.

TEs are self-replicating genomic components that are ubiquitous across eukaryotes and major drivers of genome evolution^10,11^. TEs comprise DNA transposons and retrotransposons (including LTR, non-LTR and DIRS elements), defined by their distinct transposition mechanisms and domain composition^12^. Because TE mobilization can disrupt genes and compromise genome stability, eukaryotes have evolved diverse mechanisms to regulate TE expression and propagation, including small RNA pathways, histone modifications^13–15^ and 5mC^1^. The prevalence and relative importance of these mechanisms vary widely across taxa and TE class, reflecting dynamic co-evolutionary interactions between host genomes and their TEs^2,14^. In response, TEs have evolved multiple strategies to evade or exploit host epigenetic regulation. Some TEs encode proteins that directly counter host silencing, such as plant VANDAL transposons that actively demethylate 5mC within their own sequences^15,16^. Others minimize deleterious effects by targeting insertions to permissive chromatin environments; for example, chromovirus retrotransposons use chromodomains to recognize heterochromatic histone marks and avoid integration within highly transcribed genes^17,18^. In parallel, 5mC itself has been repeatedly co-opted by TEs, with several retroelements acquiring cytosine DNA methyltransferase domains (DNMTs)^19^, TET demethylases^20^ or methyl-CpG binding proteins^21^ that may modulate local chromatin states, direct insertion sites or evade host detection. These observations highlight the importance of acquiring epigenetic-associated domains in the co-evolutionary dynamics between TEs and their hosts. However, how TEs recruit or repurpose DNA methylation systems to promote their own propagation remains poorly understood, particularly in the context of 6mA.

Intriguingly, several eukaryotic DIRS (Dictyostelium Intermediate Repeat Sequence) retrotransposons encode DAM-like N6-adenine DNA methyltransferases^22,23^. DAM enzymes are widespread in bacteria and bacteriophages, where they function as motif-specific methyltransferases involved in restriction–modification systems, DNA replication, and phage infection cycles^24,25^. While canonical DAM methyltransferases are not typically encoded by eukaryotic nuclear genomes, they have been identified in nucleocytoplasmic large DNA viruses infecting eukaryotes, which have 6mA in their genomes^26^. By contrast, the primary eukaryotic 6mA methyltransferases, including the AMT1 family mentioned above, belong to the MT-A70 superfamily and are structurally unrelated to bacterial DAM enzymes^27^.

These observations raise several unresolved questions. First, the evolutionary origin of DAM methyltransferases in eukaryotic retrotransposons is unclear, as it does their distribution beyond DIRS elements. Second, their potential functional relationship with endogenous 6mA pathways and whether TE-encoded DAM enzymes interact with host adenine methylation systems remains unknown.

To address these questions, we traced the evolution and genomic context of DAM methyltransferases encoded by eukaryotic TEs. Across diverse eukaryotes, we identified DAM domains in DIRS elements, but also recurrent co-options into LTR retrotransposons. Whereas DIRS-associated DAMs show no consistent relationship with endogenous 6mA, LTR ones are found in taxa with endogenous 6mA, and are enriched for adenine-methylated insertions. These findings suggest that prokaryotic-like methyltransferases have been independently recruited by retrotransposons and might serve to exploit 6mA as an evolutionary advantageous strategy.

## RESULTS

### Landscape of DAM-containing proteins in eukaryotes

To identify DAM-like methyltransferases in eukaryotes, we searched for DAM domains across a comprehensive collection of genomes and transcriptomes from eukaryotes (Supplementary Fig. 1A; see Methods). This analysis recovered 10,030 DAM-containing proteins with a widespread distribution across the eukaryotic tree (Fig. 1A). Phylogenetic reconstruction including representative archaeal, bacterial and viral DAM sequences from InterPro^28^ revealed that the vast majority of eukaryotic sequences form a well-supported monophyletic clade (bootstrap support = 100%; Fig. 1B). This topology suggests that most eukaryotic DAM domains derive from a single ancestral acquisition from prokaryotes. We identified five notable exceptions with more restricted phylogenetic distributions, corresponding to three putative non-viral horizontal gene transfer events and two groups associated with eukaryotic viruses.

**Figure 1.**
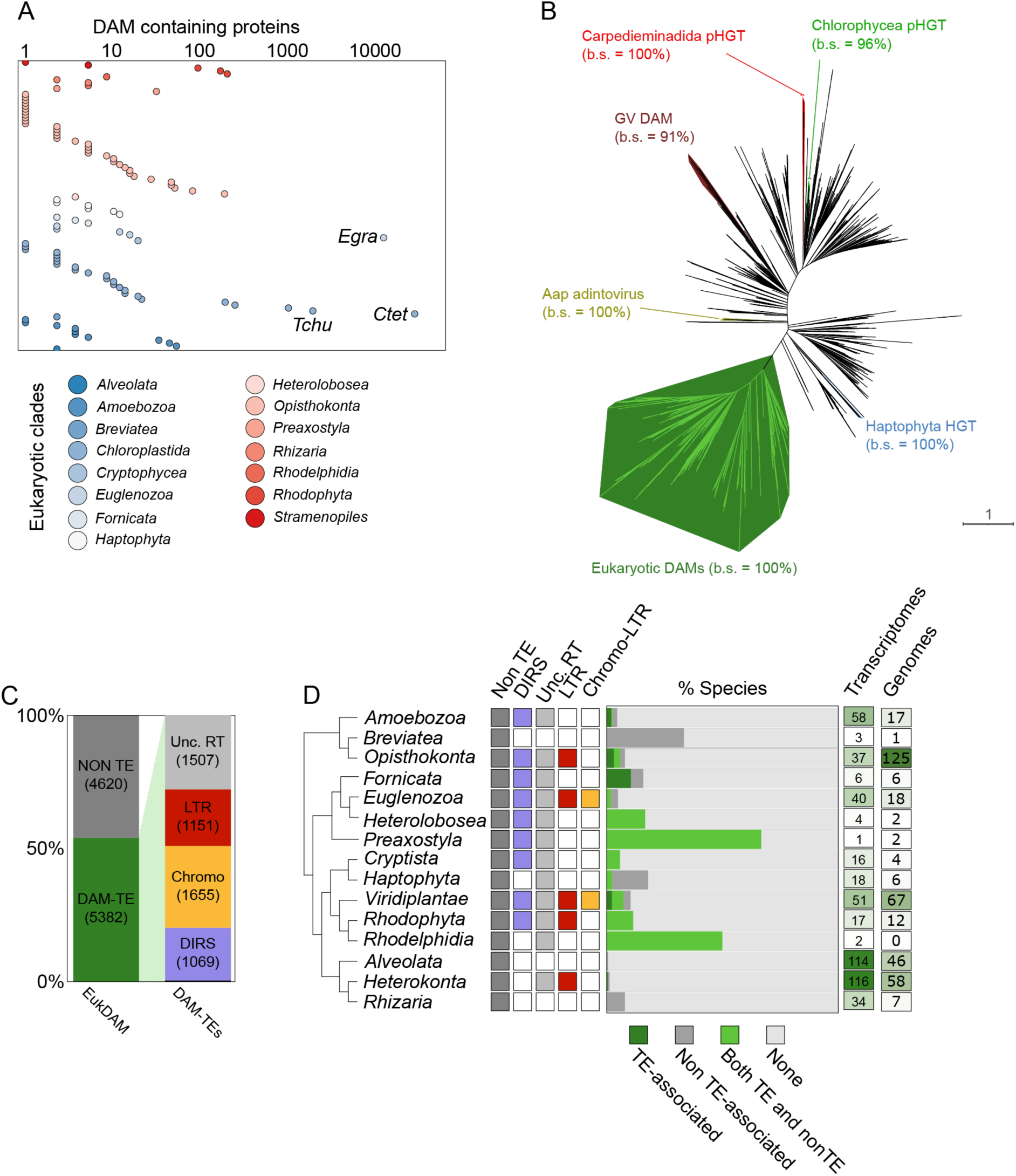
TE-associated DAM methyltransferases present a widespread distribution along eukaryotic phylogeny. A) Number of DAM-containing proteins identified per species, coloured by eukaryotic supergroup. B) Maximum likelihood phylogenetic tree of DAM domains across prokaryotes (in black) and eukaryotes. Each eukaryotic clade with more than 5 sequences is highlighted in different colours. The main shaded clade belongs to the Dam-retrotransposon clade. C) Percentage of eukaryotic DAM proteins associated with TE domains and TE classes containing DAM domains. D) Summary of the DAM screening, presence and absence of DAM-TEs by class and by taxon, percentage of species containing DAM proteins, and sample size by taxon.

The three putative non-viral horizontal gene transfer (HGT) events comprise distinct clades in Chlorophyta, Carpediemonadida, and a mixed Haptophyta–Heterokonta group, each showing affinity to different prokaryotic donors relative to the main eukaryotic DAM clade. The Chlorophyta-derived family includes multiple DAM copies in six Chlamydomonadales species (bootstrap support, b.s. = 96%) and clusters with a bacterial sequence from *Armatimonadota* (b.s. = 81%). The Carpediemonadida clade (b.s. = 100%) contains proteins from *Carpediemonas membranifera*, *Dysnectes brevis* and *Kipferlia bialata*. The Haptophyta-associated group (b.s. = 100%) comprises sequences from five Prymnesiophyceae species together with one sequence from the heterokont *Florenciella* sp., suggesting a shared acquisition.

In the case of the two groups of virus-associated DAMs, they correspond to adintoviruses and giant viruses, both known to endogenize in eukaryotic genomes. Adintovirus-related DAMs were detected in the ichthyosporean *Amoebidium appalachense*. Giant virus-like DAMs were found in transcriptomes from Amoebozoa (*Cavostelium apophysatum*, *Soliformovum irregulare*), Euglenozoa (*Peranema trichophorum*, *Rhabdomonas costata*), Chlorophyta (*Polyblepharides amylifera*) and Foraminifera (*Bolivina argentea*), as well as in genomes from the amoebozoan *Vermamoeba vermiformis*, the euglenozoan *Euglena gracilis* and the filasterean *Pigoraptor chileana*, consistent with viral integration. Notably, a DAM-encoding giant virus infecting *Vermamoeba vermiformis* has been previously reported^26^.

Despite these additional HGT events, the vast majority of eukaryotic DAM sequences cluster within a single well-supported clade, consistent with a common origin. We therefore refer to this group as the eukaryotic DAM family throughout this work. This distribution could reflect an ancient acquisition of a prokaryotic DAM by a retrotransposon early in eukaryotic evolution.

### DAM-containing proteins are associated with retrotransposons in eukaryotes

Beyond viral sequences, DAM-like domains in eukaryotes have previously been reported only in a subset of DIRS retrotransposons. Furthermore, the uneven distribution of eukaryotic DAM genes, ranging from a few copies in some species to thousands in distantly related lineages (Fig. 1A), is also suggestive of TE–mediated amplification. To test this, we assessed the association between eukaryotic DAM genes and TEs using a domain-based classification pipeline that identifies TE-related proteins within the same ORF or in their immediate genomic vicinity (see Methods; Supplementary Fig. 1A).

Using this approach, 53.9% of eukaryotic DAM proteins were associated with retrotransposon features, including reverse transcriptase (RVT-1) domains or the phage-like tyrosine integrase characteristic of DIRS elements (Fig. 1C; Supplementary Fig. 1B). We collectively refer to these as DAM-TEs. Among them, LTR retrotransposons represented the most abundant classes, with chromodomain-containing and non-chromodomain LTR elements accounting for 30.9% and 21.3% of DAM-TEs, respectively. To our knowledge, this represents the first identification of DNA adenine methyltransferase domains encoded within (non-DIRS) LTR retrotransposons (hereafter DAM-LTRs). In contrast, DIRS-like elements comprised 19.8% of DAM-TEs but displayed a broader phylogenetic distribution across eukaryotes (Fig. 1D). This proportion is likely underestimated, as DAM and the characteristic DIRS tyrosine integrase are typically encoded in separate ORFs, which limits detection to chromosome-scale assemblies or long scaffolds^29^.

Annotation limitations, including TE fragmentation, incomplete assemblies, and nested insertions, likely account for the remaining 46.1% of DAM sequences lacking detectable TE features. Consistent with this interpretation, most of these orphan DAM-only sequences were closely associated with DAM-TE branches in the phylogenetic tree, and ancestral state reconstruction across the main eukaryotic DAM clade supports a TE-encoded origin (Supplementary Fig. 1C). This indicates that most orphan DAMs likely represent degenerated TE-derived remnants rather than endogenous members of an unrelated eukaryotic gene family. This reconstruction further suggests that the initial acquisition of a prokaryotic DAM by eukaryotes was mediated by a transposable element.

At a quantitative level, eukaryotic DAM proteins display a highly heterogeneous distribution, with some species encoding hundreds to thousands of copies, whereas others contain only a few (Fig. 1A; Supplementary Fig. 1C). Comparisons across lineages indicate that these patterns are influenced by taxon sampling, with DAM-TEs detected in most eukaryotic supergroups that are represented by numerous genomes or transcriptomes (Fig. 1D), while its absence in some groups might be attributable to lack of data. Despite this broad distribution, DAM-TEs are absent from the majority of species even within deeply sampled clades, indicating that both DAM-DIRS and, in particular, DAM-LTR elements are relatively uncommon but widely dispersed and have undergone substantial expansion in a limited number of unrelated lineages.

### Recurrent evolution of DAM-LTR retrotransposons

To further investigate the evolution of DAM-TEs, we reconstructed a phylogeny of reverse transcriptase (RT; RVT1, PF00078) domains from DAM-TEs together with annotated RT sequences from Repbase (Fig. 2A). Because RT domains provide a robust phylogenetic signal for retrotransposon classification^30^, this tree was used to assign DAM-TEs to established TE families. Comparison of the RT phylogeny with the DAM-domain tree (Fig. 1B) revealed a striking contrast: whereas eukaryotic DAM domains form a largely monophyletic clade, their associated RT domains are distributed across multiple retrotransposon families. As expected, many DAM-TEs grouped within DIRS-like elements, but others branched within Ty3/Gypsy-like LTR retrotransposons. This incongruence between DAM and RT phylogenies is consistent with modular TE evolution and suggests repeated capture of a DIRS-derived DAM domain by distinct LTR retrotransposons.

**Figure 2.**
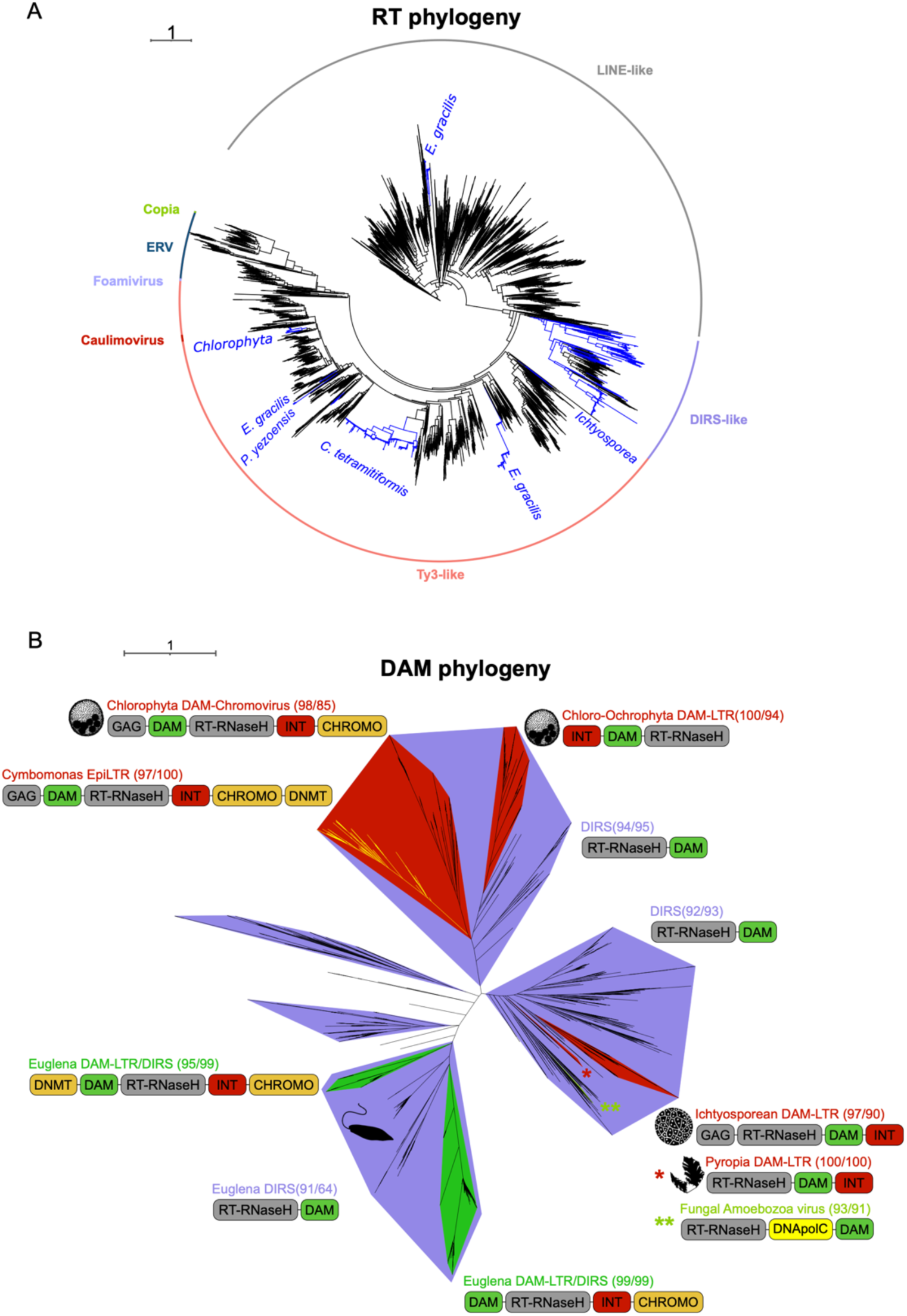
DAM-LTRs evolved convergently several times. a) Maximum likelihood phylogenetic tree encompassing reference RT domains from RepBase (black) and RTs encoded in DAM-TEs (blue). b) Maximum likelihood phylogenetic tree of DAM domains from DAM-TEs classified as LTRs (red) or DIRS (blue) by domain composition and RT phylogenetic position, also displaying the representative domain architectures. *Euglena gracilis* branches with a mixed composition of DIRS-like and LTR-like elements are labeled in green. Scale represents substitutions per site.

To examine this pattern in more detail, we reconstructed a focused phylogeny of DAM domains restricted to DAM-TEs and annotated each sequence using the RT-based classification as well as a domain co-occurrence one (Fig. 2B; Supplementary Fig. 2A-C). Most DAM domains could be confidently assigned to either DIRS or LTR retrotransposons. The tree topology supports at least four independent transfers of DAM domains from DIRS-like elements into LTR retrotransposons, plus an additional capture associated with a virus-like element encoding a hepadnavirus-like DNA polymerase. Based on these relationships, we define five independent DAM-LTR lineages.

The **Chlorophyta DAM-chromovirus-like** clade (b.s. = 77%) includes elements from multiple green algae and is closely related to DIRS-like sequences from algae and opisthokonts (b.s. = 94%). These elements typically show a chromovirus-like architecture, including a chromodomain histone modification reader. The **Chloro–Ochrophyta DAM-LTR** clade (b.s. = 100%) includes elements from chlorophytes and an ochrophyte, and appears related to the Chlorophyta DAM-chromovirus in the DAM tree but not in the RT phylogeny, consistent with independent capture events. These LTRs contain the DAM domain flanked by the integrase domain and the RT. The **Ichthyosporean DAM-LTR** lineage (b.s. = 97%), identified in *Pirum gemmata*, *Abeoforma whisleri* and *Sphaeroforma arctica*, lacks chromodomains and displays a conventional Ty3/Gypsy-like organization. The **Pyropia DAM-LTR** clade (b.s. = 100%) comprises a small number of insertions in the rhodophyte *Pyropia yezoensis* and is closely related to the same DIRS lineage as the ichthyosporean elements. Finally, the **Euglena DAM-LTR/DIRS** group consists of mixed DIRS and LTR elements forming two strongly supported clades (b.s. = 99%), consistent with recent domain exchange or nested insertion events. Notably, *Euglena gracilis* shows extensive RT diversity, including DAM-LTRs branching with LINE/L1-like RTs despite retaining a Ty3-like structure, suggesting ongoing domain transfer among retroelements in this lineage, potential nested insertions creating chimeric TEs or assembly artifacts.

### EpiLTRs: dual methylation retrotransposons

Within two of the DAM-LTR lineages described above, we identified retrotransposons encoding an unusually complex combination of epigenetic domains. In the **Chlorophyta DAM-chromovirus-like** clade, a monophyletic subset of *Cymbomonas tetramitiformis* elements displays a canonical chromovirus-like architecture but with an additional cytosine DNA methyltransferase (DNMT) appended to the 3′ end of the *pol* ORF. A similar yet structurally distinct configuration was observed in the **Euglena DAM-LTR**, where a subset of LTR retrotransposons encodes a chromodomain together with both DAM and DNMT domains. The different domain organizations and distant positions in the RT and DAM phylogenies suggest that in *C. tetramitiformis* and *E. gracilis* these complex TE architectures assembled independently.

While the convergent acquisition of DNMTs by divergent retrotransposons has been reported previously, the combined presence of two DNA methyltransferases (one associated with 6mA and the other with 5mC) together with a chromodomain has not been described previously. We refer to these elements as **EpiLTRs**, reflecting their expanded epigenetic repertoire. To further investigate their origin, we reconstructed a phylogeny of DNMT domains including representative eukaryotic DNMT families and EpiLTR-associated sequences (Supplementary Fig. 2D). DNMTs from *E. gracilis* and *C. tetramitiformis* EpiLTRs form a well-supported monophyletic group (b.s. = 100%) sister to the DNMT4 clade, suggesting a shared origin of the DNMT module, despite the RT and DAM domains indicating independent origins for the remaining retrotransposon components. This DNMT acquisition is distinct from previously described TE-associated DNMTs in Symbiodiniaceae dinoflagellates (SymbioDIRS-DNMT3 and SymbioLINE-DNMT) and streptophytes (Gypsy-DNMT3), although it mirrors the convergent recruitment of DNMT3 by unrelated DIRS and Gypsy elements in distantly related lineages.

These results identify independently assembled retrotransposons carrying both 6mA and 5mC methyltransferases alongside a Chromo chromatin-targeting domain, highlighting the modular evolution of TEs and their capacity to assemble unexpectedly complex epigenetic toolkits.

### Eukaryotic DAM domains preserve the catalytic pocket

To further understand the potential functions of the DAM domain in DAM-TEs, we predicted enzymatic activity using the EnzBert E.C. tool. In both viral and TE DAM domains the most likely activity assigned was *site-specific DNA-methyltransferase (adenine-specific)* (EC 2.1.1.72) (Fig. 3A).

**Figure 3.**
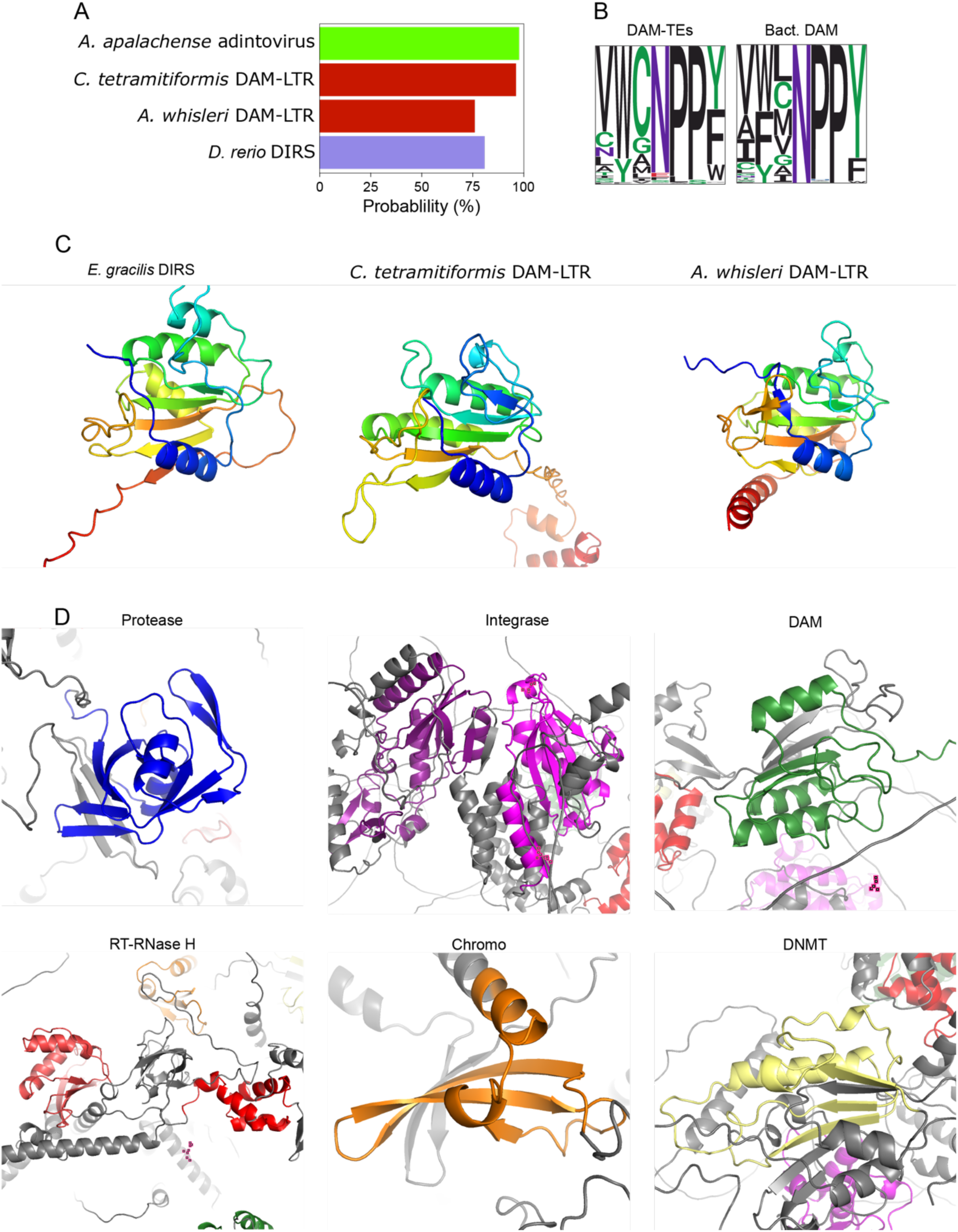
Eukaryotic DAM domain potential activity and conservation. **a**) Probability of *site-specific DNA-methyltransferase (adenine-specific)* assigned to most complete and repeated insertions of representative TE families with eukaryotic DAMs. **b**) Alignment logo of the key region of the enzymatic pocket in DAM-TEs and bacteria DAM. c) 3D prediction of representative DAM domains from TEs. d) 3D prediction of the domains contained in the polyprotein of *C. tetramitiformis* EpiLTR.

In prokaryotes, the DAM domain is split into two catalytic domains (CD-1 and CD-2) flanking the central target recognition site (TR), which itself is divided into TR-1 and TR-2 by a conserved hairpin^31^. Eukaryotic DAMs align only with prokaryotic DAMs from TR-2 to CD-2 (supplementary Fig. 3A), missing CD-1 and TR-1. Additionally, the hairpin does not appear to be conserved. Despite this sequence divergence, the key motif of the active site is well conserved (Fig. 3B).

Most of what we know about this motif comes from detailed studies of T4 phage and *E. coli*^31,32^. However, our broader taxon sampling in both eukaryotes and prokaryotes shows some differences in terms of domain architecture, providing a more complete picture. The most conserved part of this motif consists of a triplet at the catalytic pocket that in almost all sampled prokaryotes and eukaryotes DAMs is composed of Asn-Pro-Pro (Fig. 3B). Regarding the residues flanking this triplet, there are differences between the residues most commonly found in our sampled DAM and those that have been functionally characterized in T4 and *E. coli*. The initial Val (four residues upstream the triplet) is mostly conserved, but is frequently substituted by other non-polar residues, such as Ala, Ile or Leu. Both Tyr following the initial Val and the one following the Asn-Pro-Pro triplet are mostly conserved, but commonly replaced by another aromatin amino acid. Thus, the most conserved region of the DAM domain within the enzymatic pocket in prokaryotes is also well conserved across eukaryotic DAMs.

We also studied the 3D structure of the whole eukaryotic DAM domain (Fig. 3C). Despite presenting slight differences in the predicted folding, the studied domain from DIRS and DAM-LTR present a conserved structure, characterized by five strand β-sheet surrounded by an α-helix in N-terminal and 3 at the C-terminal.

Finally, we assess the viability of the EpiLTR polyprotein folding (Fig. 3D), confirming their fold into stable structures, even if post-translational cleavage may make these predictions inaccurate.

Finally, to test whether eukaryotic DAMs can catalyse adenine methylation, we expressed representative proteins from DIRS elements, DAM-LTRs, and an endogenized adintovirus in a DAM-deficient *E. coli* strain (see Methods). Genomic DNA from these strains was sequenced using Oxford Nanopore to quantify 6mA levels. While we detected the expected difference between wild-type and DAM-deficient *E. coli*, expression of the heterologous eukaryotic DAMs did not increase global 6mA levels (Supplementary Fig. 3B,C), consistent with previous heterologous expression experiments using caulimovirus DAM^33^. This negative result may reflect experimental limitations, such as protein misfolding in *E. coli*, inefficient DNA binding, or the possibility that these enzymes require a DNA–RNA hybrid substrate, as occurs during retrotransposon replication. These findings suggest that eukaryotic DAMs retain catalytic potential for 6mA deposition, although functional activity may require expression within the full retrotransposon context.

### DAM-LTRs are enriched for 6mA in AMT1-encoding eukaryotes

The presence of a DAM methyltransferase in DAM-TEs might be a co-evolutionary mechanism to adapt to host epigenomes. Therefore, we assessed the co-occurrence of DAM-DIRS elements and DAM-LTRs with AMT1, the primary eukaryotic 6mA methyltransferase, whose presence is the best predictor of abundant 6mA at ApT dinucleotides associated with transcriptionally permissive chromatin across eukaryotes. We constructed a phylogenetic tree of eukaryotic adenine methyltransferases, incorporating representative taxa and the 48 species in which complete DIRS and DAM-LTRs were identified (Fig. 4A). DIRS elements did not show a clear association with AMT1, being especially prevalent in Dyctiostelea and Metazoa, clades where loss of both 6mA and AMT1 has been previously reported. By contrast, DAM-LTRs were predominantly found in species encoding AMT1 (9 of 13 DAM-LTR-positive species were also AMT1 positive in our screen; *P* = 1.5e-04, Chi-squared test). Notable exceptions included the rhodophyte *P. yezonensis* and the chlorophytes *C. tetraminiformis (*BUSCO score 74.6%^34^)*, D. salina* (BUSCO score 89%^35^), and *O. quekettii* (BUSCO score 60.7%^36^). Given the presence of AMT1 in most chlorophytes, we hypothesize that the absence of AMT1 in these three chlorophytes may be caused by genome assembly or sequencing biases. The association between AMT1 and DAM-LTRs is further supported by the absence of these TEs in Metazoa and Streptophyta—both lacking abundant genomic 6mA—despite their presence in sister groups (Ichthyosporea and Chlorophyta, respectively).

**Figure 4.**
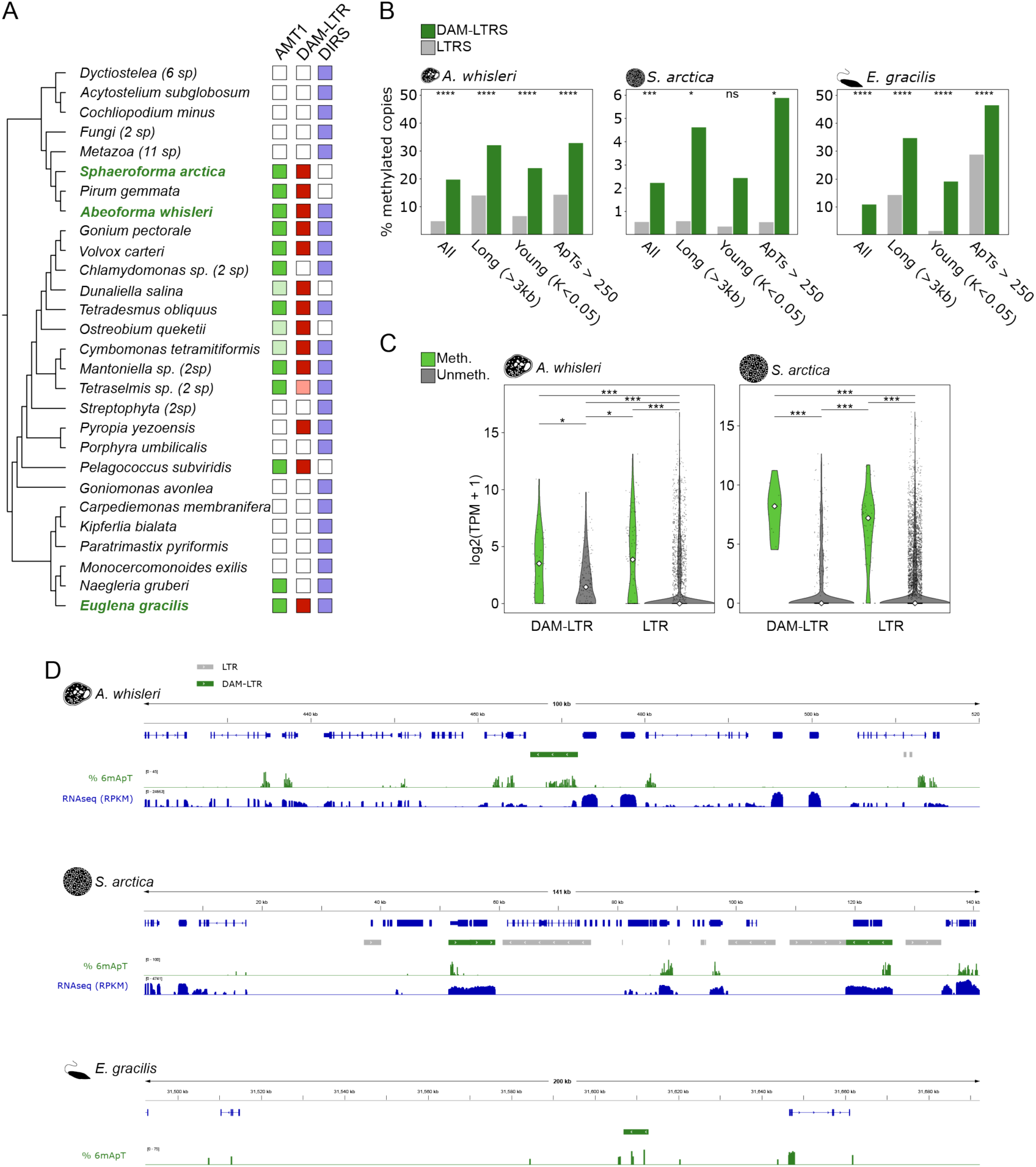
DAM-TE adenine methylation and transcriptional activity. a) Phylogenetic distribution of AMT1 and DAM-TE classes across the eukaryotes. White boxes indicate absence, while coloured boxes indicate presence. Pale colors reflect partial evidence of AMT1 or DAM-LTR presence. b) Percentage of methylated insertions in DAM-LTRs and DAM-lacking LTRs in *A. whisleri* and *S. arctica* establishing increasing thresholds for considering the insertion as methylates. c) Normalized expression of DAM-LTR and DAM-lacking LTRs in *A. whisleri* and *S. arctica*. d) Representative genome browser captures of regions with DAM-LTRs in *A. whisleri, S. arctica* and *E. gracilis* showing RNAseq normalized signal percentage of 6mApT. Repeats are shown in grey, and DAM-LTRs are shown in green.

To investigate the functional relationship between DAM-LTRs, 6mA methylation, and transcription, we analyzed the TE 6mA profiles in two DAM-LTR–containing ichthyosporeans, using previously published data for *Abeoforma whisleri* and newly generated data for *Sphaeroforma arctica* (Fig. 4A–D; Supplementary Fig. 4A and 5B). We profiled the *S. arctica* 6mA methylome, which displayed the hallmark characteristics of 6mA in ichthyosporeans, accumulating after the TSS of transcribed genes, displaying a periodical pattern presumed to be internucleosomal (Supplementary Fig. 4B). Interestingly, although 6mA is generally enriched in endogenous genes and absent in TEs, we detected a significant increase in the proportion of methylated DAM-LTR insertions relative to other LTRs (Fig. 4B). Importantly, this enrichment persisted regardless of the threshold used to define methylated insertions, using TE length or predicted age (Kimura distance), demonstrating that it is specifically associated with a specific LTR type (the DAM-LTRs) and not simply with other LTR features such as completeness or age.

Beyond ichthyosporeans, we also examined the euglenozoan *E. gracilis* (Supplementary Fig. 4B). Its epigenome is markedly more complex than that of ichthyosporeans, with 5mC being almost the default state at CG dinucleotides, including across most gene promoters. In contrast, 6mA is enriched downstream of the TSS, but only on a small subset of genes whose promoters strikingly lack CpG methylation. This configuration represents a highly divergent epigenomic architecture compared with other eukaryotes and suggests the existence of two promoter classes: one resembling the canonical state (low 5mC and detectable 6mA) and another that is heavily methylated at CpG sites. The prevalence of promoter methylation in *E. gracilis* may be associated with its extensive trans-splicing^37,38^. This is a feature also associated with promoter methylation in two distantly related lineages that use trans-splicing: the ctenophore *Mnemiopsis leidyi* and symbiodiniacean dinoflagellates^19,39^. In line with our results in ichthyosporeans, *E. gracilis* also showed an increased proportion of DAM-LTRs methylated (Fig 4B and D). This increase in both distant clades whose DAM-LTRs evolved independently reinforce the importance of the DAM domain for these new LTR classes.

Consistent with the presence of 6mA, we also detected an increased expression of DAM-LTRs in both *S. arctica* and *A. whisleri* (Supplementary Fig. 5C), which was positively associated with transcriptional activity of these TEs (Fig. 4C). This tendency was not observed in *E. gracilis*, an organism in which the general expression of LTRs was low or undetectable (supplementary Fig. 5D). The concomitant increase in 6mA and expression observed in DAM-containing LTRs suggests a functional link between these features.

## DISCUSSION

We show that retrotransposons encoding prokaryotic-like DAM methyltransferases are widespread across eukaryotes and derive from a largely monophyletic eukaryotic DAM lineage. The phylogenetic clustering and broad taxonomic distribution support an ancestral acquisition of a prokaryotic-like DAM domain by a DIRS-like retrotransposon early in eukaryotic evolution, followed by vertical inheritance and lineage-specific expansions. The presence of phage-like components in DIRS elements, including the tyrosine recombinase and DAM domains, further supports a potential origin involving viral or phage-related donors. Thus, DAM-encoding DIRS-like elements are likely part of the list of extant retrotransposons already present in the Last Eukaryotic Common Ancestor (LECA)^40^.

Our analyses indicate that DAM domains, that were originally found in these DIRS-like elements, were later repeatedly transferred into Ty3/Gypsy-like LTR retrotransposons across different eukaryotes. We identify at least four independent DAM-LTR lineages, with *Euglena gracilis* displaying a more complex scenario consistent with multiple exchanges or nested insertions. The recurrent emergence of DAM-LTRs suggests that retention of the DAM domain is functionally advantageous for the TEs. Notably, most DAM-LTR lineages occur in species encoding AMT1, whereas DIRS elements show no such association and are abundant in lineages lacking 6mA. This contrast suggests that recruitment of DAM by LTR retrotransposons involved a functional shift, potentially adapting these elements to epigenomes where 6mA marks transcriptionally permissive chromatin.

Consistent with this interpretation, DAM-LTR insertions show increased 6mA methylation relative to other LTR elements. Although 6mA is generally depleted from TEs^6^. DAM-LTRs display a higher proportion of methylated copies, particularly among younger and more complete insertions. This enrichment suggests that TE-encoded DAMs may facilitate deposition of 6mA, either directly or by promoting recruitment of host AMT1. One possible model is that DAM activity deposits 6mA on reverse-transcribed RNA–DNA intermediates, which upon integration recruit host AMT1, thereby maintaining 6mA and a permissive chromatin environment for new insertions. Elevated expression of DAM-LTRs in ichthyosporeans further supports a link between 6mA and transcriptional permissiveness, although this relationship may depend on lineage-specific epigenomic contexts, as illustrated by the low overall LTR expression in *E. gracilis*. While methylation data are currently limited to a subset of species with active DAM-LTR, their independent origins suggest a shared functional advantage. Together, these observations support a potential strategy to exploit host epigenetic regulation.

We also identify a subset of retrotransposons combining multiple epigenetic modules, including chromodomains together with both 6mA and 5mC methyltransferases. These EpiLTRs highlight the modular evolution of TEs and their capacity to assemble complex epigenetic toolkits, in some cases encoding a broader repertoire of DNA methylation activities than those found in model eukaryotes such as yeast or flies. Why such a combination of epigenetic regulators is maintained within a single retrotransposon family remains unclear, but it underscores the importance of epigenome interaction as a key adaptive barrier that TEs must navigate to persist within host genomes. *Euglena* EpiLTRs are a particularly intriguing example, as the *Euglena* epigenome is heavily methylated in 5mC while also containing promoter-associated 6mA at subsets of endogenous genes, suggesting that the ability to engage with both epigenetic systems may provide a selective advantage in this context.

The recurrent co-option of methyltransferases and chromatin-binding domains indicates that TE–host interactions frequently operate at the epigenomic level, although the outcome of these acquisitions likely depends on the regulatory landscape of each host. In some cases, these domains may enhance TE transcription, such as TET or VANDAL-associated 5mC demethylases^15,16^. In others, they may minimize deleterious insertions through co-evolutionary strategies, as exemplified by chromodomains that guide integration into gene-poor regions^41^. In this context, 6mA-associated chromatin appears particularly amenable to TE exploitation in lineages that retain this modification, as it can be co-opted to promote transcriptionally permissive chromatin. More broadly, these findings reinforce the view of the genome as an evolving ecosystem in which TEs continuously acquire and redeploy regulatory modules to persist within dynamic host environments. In turn, acquisition of prokaryotic DNA methyltransferases through lateral gene transfer can shape eukaryotic epigenomes, as exemplified for 4-methylcytosine in rotifers and the liverworts^42,43^. As host epigenomes diversify and counter-adapt, these interactions generate a shifting evolutionary landscape, in which both TEs and host genomes must continually evolve to maintain their relative fitness.

## METHODS

### Eukaryotic DAM screening and DAM-TE classification

Eukaryotic DAMs were identified from a subset of 880 reference genomes (Supplementary Table 1) and transcriptomes (Supplementary Table 2) available in EukProt V3^44^. Genomes derived from single-cell or bulk DNA-seq were translated into open reading frames using getORF (EMBOSS v6.6.0.0^45^) to minimize false positives arising from misannotated splice junctions. Taxonomic assignment was based on EukProt V3 metadata, and the reference phylogeny followed Williamson et al.^46^. Proteomes derived from single-cell and bulk RNA-seq were used directly. DAM-containing proteins and their domain architectures were identified using *hmmsearch* from HMMER v3.3.2^47^ applying an E-value threshold of 10^-5^. Transposable elements (TEs) were detected and classified according to their domain composition (Supplementary Fig. 1A). DIRS retroelements were identified by extracting a 3 kb window surrounding the DAM domain ORF and searching for the presence of open reading frames encoding phage-like tyrosine recombinases within this region.

### Phylogenetic analysis

For each phylogenetic analysis, multiple sequence alignments were generated using MAFFT v7 with the L-INS-i algorithm^48^. Alignments were trimmed using trimAl^49^ with the “-gappyout” option, and redundancy was removed and visualized with Jalview v2.11.5.0^50^. Phylogenetic trees were inferred with IQ-TREE2^51^ employing automatic model selection and evaluating branch support with 1,000 ultrafast bootstrap replicates, 1,000 SH-aLRT replicates, and aBayes. For the eukaryotic and prokaryotic DAM tree, all proteins containing DAM domains from Archaea, Bacteria, and Phages were downloaded from InterPro v106.0^28^ and then filtered to remove proteins with more of 95% of redundancy. For the RT tree, reverse transcriptase domains were extracted from the Repeat Protein Database^52^. In the DAM-TEs tree, DAM domain-containing sequences were labeled according to their RT tree clustering and the domain classification described above. For the DNMT tree, representative sequences were obtained from Sarre et al.^53^ The MT-A70 domain alignment was constructed using representative sequences from Romero Charria et al.^6^ and performing a tblastn (BLAST 2.12.0+) search against the genomes of species that present DAM-DIRS or DAM-LTRs.

### Domain characterization

Enzymatic activity was predicted using EnzBert E.C.^54^. Sequence logos for enzymatic pockets were generated using the R package ggseqlogo^55^.

Structural predictions were performed using AlphaFold 3^56^, and the resulting models were superimposed using PyMOL v3.

### Heterologous methylation assay in *E. coli*

Representative DAM sequences (Supplementary Table 2) were synthesized by Twist Bioscience and cloned into the pTwist Amp High Copy vector under the control of a Ptac promoter. Each insert was flanked by an *E. coli* ribosome binding site (RBS) and T1 terminator. An eGFP-encoding plasmid served as a negative control. Experiments were conducted in DAM-deficient *E. coli* One Shot™ INV110 cells, with competent DH5α cells used as a positive control. Transformations were carried out according to the manufacturer’s protocol, and protein expression was induced by adding IPTG to the culture medium. Genomic DNA was extracted using the Monarch® Spin gDNA Extraction Kit (NEB #T3017). For profiling the global 6mA methylation level in these heterologous DAM strains, multiplexed libraries were prepared using the Oxford Nanopore Rapid Barcoding Kit V14 (SQK-RBK114.96) following manufacturer’s instructions and sequenced on PromethION R10.4.1 flow cells. Pod5 raw read files were base-called, mapped, and de-multiplexed by Dorado (v0.7.2). The modifications were identified using the pileup function of Oxford Nanopore Modkit (v0.4.2), using 0.95 threshold for modified base.

6mA modifications were identified using the pileup function of Oxford Nanopore Modkit v0.4.2, evaluating multiple thresholds for modified base calling.

### *S. arctica* cell culture and DNA extraction and sequencing

*S. arctica* (JP610) was grown in Marine Broth liquid media (Difco, 2216) at 17 °C, using 25 ml flasks. A confluent culture was grown for 6 days, cells were pelleted using centrifugation at 3,000g for 5 min, and the supernatant was discarded. To break the cell wall the pellet was frozen with liquid nitrogen and thawed at 60°C, repeating this thrice. For DNA extraction NEB Monarch Genomic DNA Purification Kit was used following the animal tissue protocol.. The resulting DNA was then analysed with a TapeStation 2200 to visualize genomic DNA integrity using the Genomic DNA ScreenTape Assay and quantified using a Qubit 3 Fluorometer with the dsDNA BR Assay Kit and then high molecular weight genomic DNA was used for Oxford Nanopore ligation sequencing kit V14 (SQK-LSK114) following the manufacturer’s instructions starting with 1.5μg of DNA, and sequenced later in PromethION R10.4.1 flow cell.

### ONT epigenome characterization

Base-modified calls for *S. arctica* were generated using Dorado (v0.7.2) with the sup model to detect 6mA, these modifications were identified using the pileup function of Oxford Nanopore Modkit (v0.4.2), using 0.995 threshold for modified base as shown in Romero Charria et al. (2025). For *E. gracilis*, basecalled 6mA and 5mC data was downloaded from Zenodo https://doi.org/10.5281/zenodo.1523872357.

Modified base frequencies (6mA, 6mA at ApT, 5mC and CpG methylation) were computed genome-wide using modkit (v0.5.0). Resulting methylation tracks were converted to bigWig format in R using the rtracklayer Bioconductor package. These tracks were used as input for deepTools (v3.0)^58^ to compute metaprofiles using protein-coding genes as the reference BED file. For E. gracilis, an additional CpG density track was generated using 100 bp windows.

### *E. gracilis* gene annotation

Because the released *E. gracilis* gene annotation lacked UTRs and showed limited accuracy, we generated an improved transcriptome-based annotation. A *de novo* transcriptome assembly from a previous study^38^ was mapped to the reference genome^59^ using GMAP. Short-read RNA-seq datasets (SRR17465012, SRR17465011, SRR17465010, SRR17465022, SRR17465030) were aligned with HISAT2 using the --dta option. Each BAM file was assembled independently with StringTie3, and high-confidence splice junctions were identified using Portcullis. These transcript models were integrated with Mikado, which selected the best-supported gene model per locus using DIAMOND searches against UniProt as reference.

### TEs methylation quantification in *A. whisleri*, *S. arctica* and *E. gracilis*

For *A. whisleri* nanopore 6mA reads were downloaded from BioStudies S-BSST1363^6^. For S. arctica and E. gracilis, the modified bases frequencies generated from the epigenome characterization (see above) were used.

EarlGrey^60^ was used for TEs identification and classification. DAM-LTR families were identified as the best tblastn (BLAST 2.12.0+) hit of each previously identified DAM domain containing proteins of each specie. The rest of LTR families identified were considered as non DAM containing LTRs. Methylation levels of the LTR insertions were analyzed in R, using the Bioconductor bsseq^61^ package trating the ApT dinucleotides as CpGs. To consider a site as mehtylated, the 6mA percentage should be higher than 10% and the coverage above 10x. To consider a TE insertion as methylated, 10 methylated ApTs were required for *A. whisleri* and *E. gracilis* and 5 methylated ApTs for *S. arctica* (coherently with its intrinsically less methylated genome). The difference in amount of methylated insertions in DAM-LTRs and LTRs was tested by a χ² test using the Benjamini-Hochberg FDR procedure.

For transcriptomic analysis, we downloaded public data from RNAseq experiments of *A. whisleri* (SRA PRJNA360056) and *S. arctica* (SRA PRJNA1048300). The fastq files were aligned using STAR 2.7.10a^62^ and TE expression was quantified using featureCounts 2.0.3^63^ allowing only uniquely Mappable Reads to avoid false positive counts^64^. Counts were normalized by TPMs and differential expression was assayed by Kruskal-Wallis test using the Benjamini-Hochberg FDR procedure. Transcriptomic and 6mA levels were visualized using IGV 2.17.0^65^. For RNAseq visualization, reads were normalized by RPKM employing a 20 bp bin size.

## Acknowledgements

We would like to thank Benjamin Tapon and Peng Zhou for helpful discussions on cloning and enzyme phylogenetics, and Iñaki Ruiz-Trillo, Elena Casacuberta and Meritxell Antó for helping with *Sphaerforoma* cultures. MFM was funded by a EMBO Scientific Exchange Grant (ref. 11468) and a Spanish Ministry of Universities FPU grant (ref. FPU20/02733). IM and MFM were supported by grants PID2021-128728NB-I00 and CNS2022-136105 funded by MICIU/AEI/10.13039/501100011033 and by ‘ERDF/EU’ and ‘European Union NextGenerationEU/PRTR’. This work used computing resources from Queen Mary University of London’s Apocrita HPC facilities. This work was funded by the Horizon 2020 Framework Programme (European Research Council Starting Grant action 950230 to AdM). PRC was funded by a QMUL PhD fellowship.

## Data availability

The Nanopore data generated in this manuscript can be found in GEO under accession number GSE331346.

## Author contributions

MFM, AdM and IM designed and conceptualised the study. MFM performed the phylogenetic analyses, TE annotations, plasmid cloning and expression experiments, and analysed the 6mA data. PRC and LX generated ONT libraries and contributed to basecalling and downstream analyses together with AdM. BLV assisted with bacterial plasmid cloning. MI, AdM and NM supervised the work. MFM, AdM, NM and MI wrote the first draft of the manuscript, and all authors contributed to subsequent revisions.

