## SupplementaryFigures for "Pervasive co-option of prokaryotic adenine methyltransferases by eukaryotic retrotransposons"

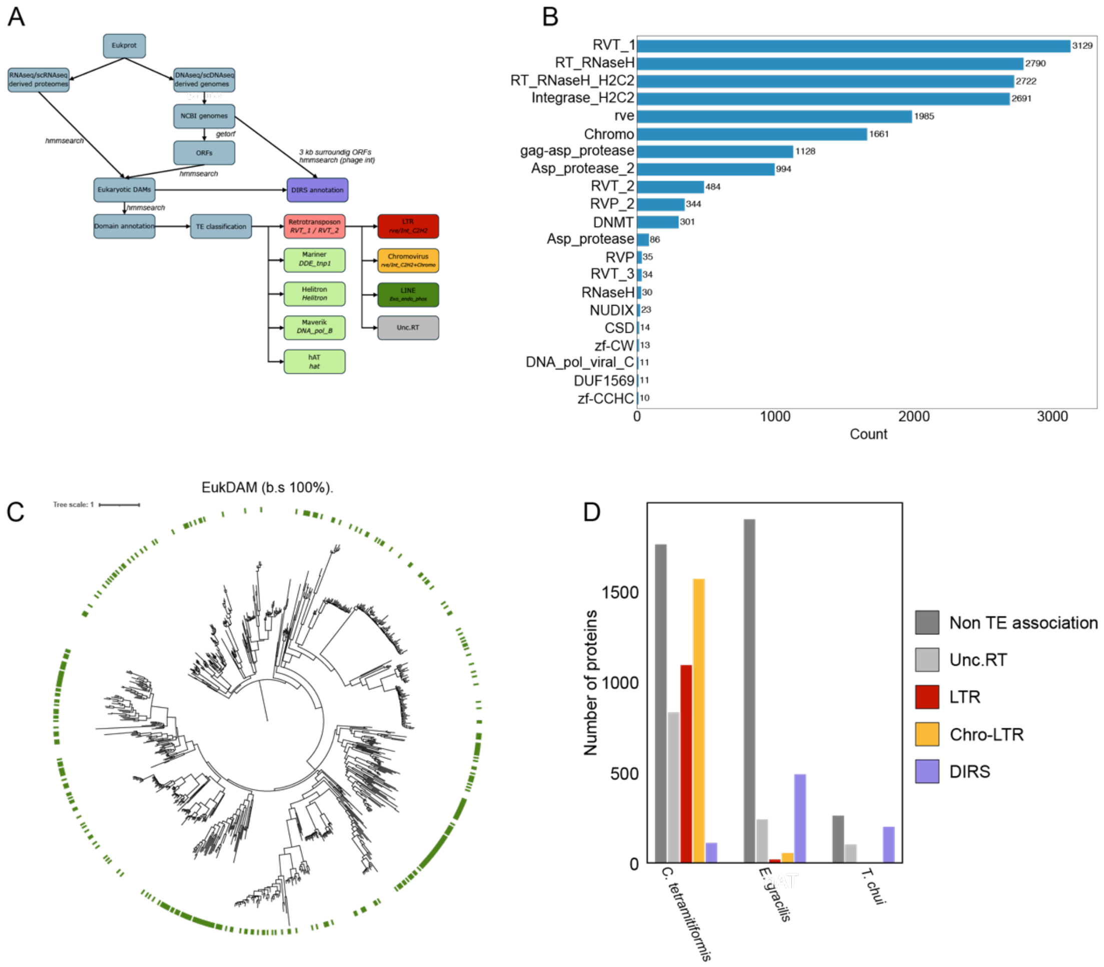


**Supplementary figure 1. Identification and classification of DAM retrotransposons.** A) Workflow used for eukaryotic DAM screening and DAM-TE classification. B) Most common concomitant domains to DAM in the same ORF. C) Pruned phylogenetic (from Fig 1B) tree of the main eukaryotic DAM branch. The green labels mark the retrotransposon-associated proteins. D) DAM-TE content of the species with highest DAM-TEs counts.

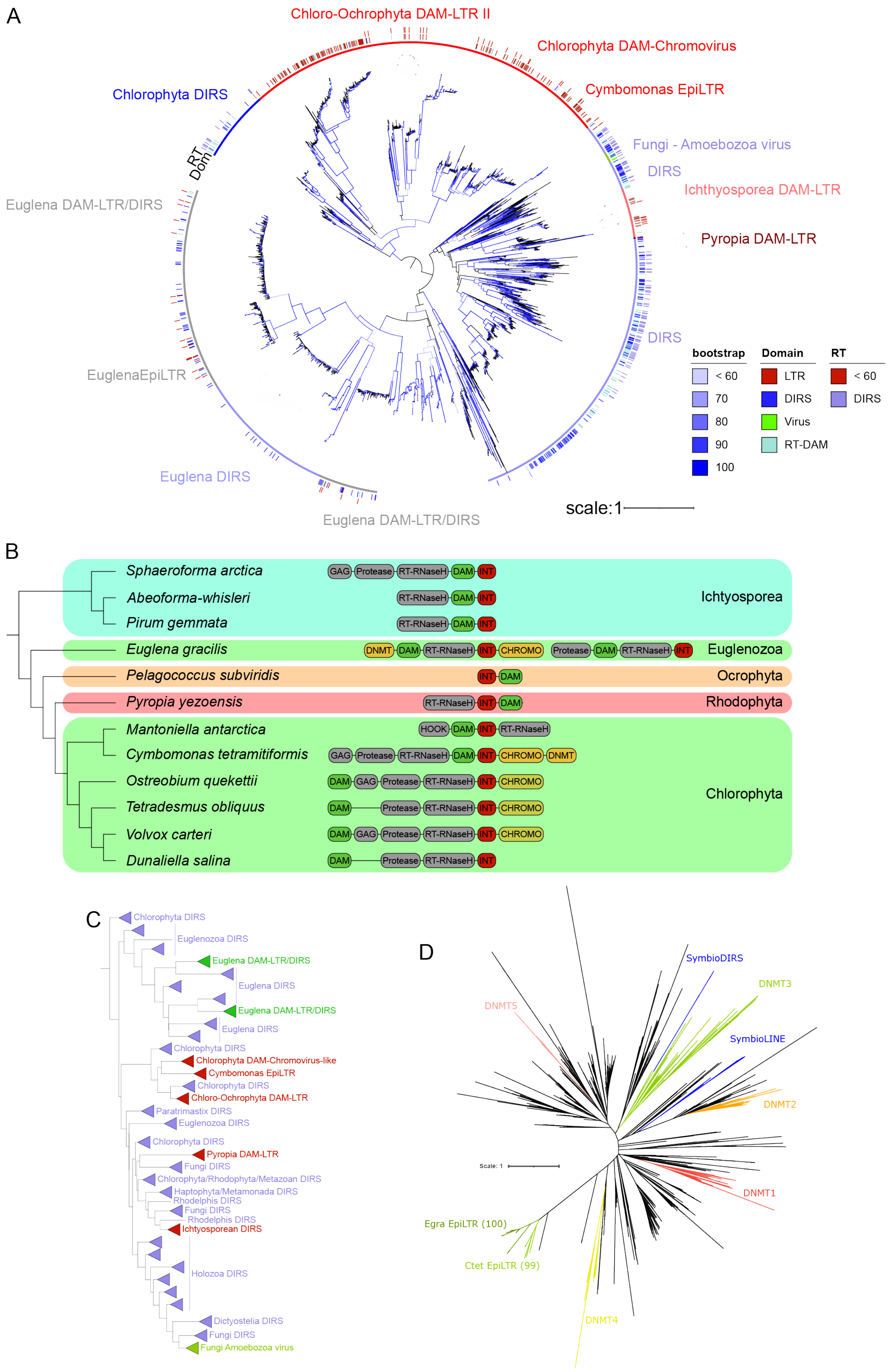
A
Chloro-Ochrophyta DAM-LTR II
Chlorophyta DAM-Chromovirus
Cymbomonas EpiLTR
Fungi - Amoebozoa virus
DIRS
Ichthyosporea DAM-LTR
Pyropia DAM-LTR
DIRS
Chlorophyta DIRS
RT
Dom
Euglena DAM-LTR/DIRS
EuglenaEpiLTR
Euglena DIRS
Euglena DAM-LTR/DIRS
bootstrap
< 60
70
80
90
100
Domain
LTR
DIRS
Virus
RT-DAM
RT
< 60
DIRS
scale:1
B
Sphaeroforma arctica
Abeoforma-whisleri
Pirum gemmata
Ichtyosporea
Euglena gracilis
Euglenozoa
Pelagococcus subviridis
Ocrophyta
Pyropia yezoensis
Rhodophyta
Mantoniella antarctica
Cymbomonas tetramitiformis
Ostreobium quekettii
Tetradesmus obliquus
Volvox carteri
Dunaliella salina
Chlorophyta
GAG Protease RT-RNaseH DAM INT
RT-RNaseH DAM INT
RT-RNaseH DAM INT
DNMT DAM RT-RNaseH INT CHROMO Protease DAM RT-RNaseH INT
INT DAM
RT-RNaseH INT DAM
HOOK DAM INT RT-RNaseH
GAG Protease RT-RNaseH DAM INT CHROMO DNMT
DAM GAG Protease RT-RNaseH INT CHROMO
DAM Protease RT-RNaseH INT CHROMO
DAM GAG Protease RT-RNaseH INT CHROMO
DAM Protease RT-RNaseH INT
C
Chlorophyta DIRS
Euglenozoa DIRS
Euglena DAM-LTR/DIRS
Euglena DIRS
Euglena DAM-LTR/DIRS
Euglena DIRS
Chlorophyta DIRS
Chlorophyta DAM-Chromovirus-like
Cymbomonas EpiLTR
Chlorophyta DIRS
Chloro-Ochrophyta DAM-LTR
Paratrimastix DIRS
Euglenozoa DIRS
Chlorophyta DIRS
Pyropia DAM-LTR
Fungi DIRS
Chlorophyta/Rhodophyta/Metazoan DIRS
Haptophyta/Metamonada DIRS
Rhodelphis DIRS
Fungi DIRS
Rhodelphis DIRS
Ichtyosporean DIRS
Holozoa DIRS
Dictyostelia DIRS
Fungi DIRS
Fungi Amoebozoa virus
D
SymbioDIRS
DNMT3
DNMT5
SymbioLINE
DNMT2
Scale: 1
DNMT1
Egra EpiLTR (100)
Ctet EpiLTR (99)
DNMT4

**Supplementary figure 2. Phylogenetic relationships and domain architectures of DAM containing retrotransposons.** a) Detailed maximum likelihood phylogenetic tree of DAM domains from DAM-TEs classified by RT phylogeny and domain architecture. b) Representative domain architectures of the DAM-LTRs found in various species. c) Simplified phylogenetic tree of the DAM-TE DAM domain and their classification. d) Maximum likelihood phylogenetic tree of DNMT domain from representative species and the *C. tetramitiformis* (Ctet) and *E. gracilis* (Egra) LTR-associated DNMT.

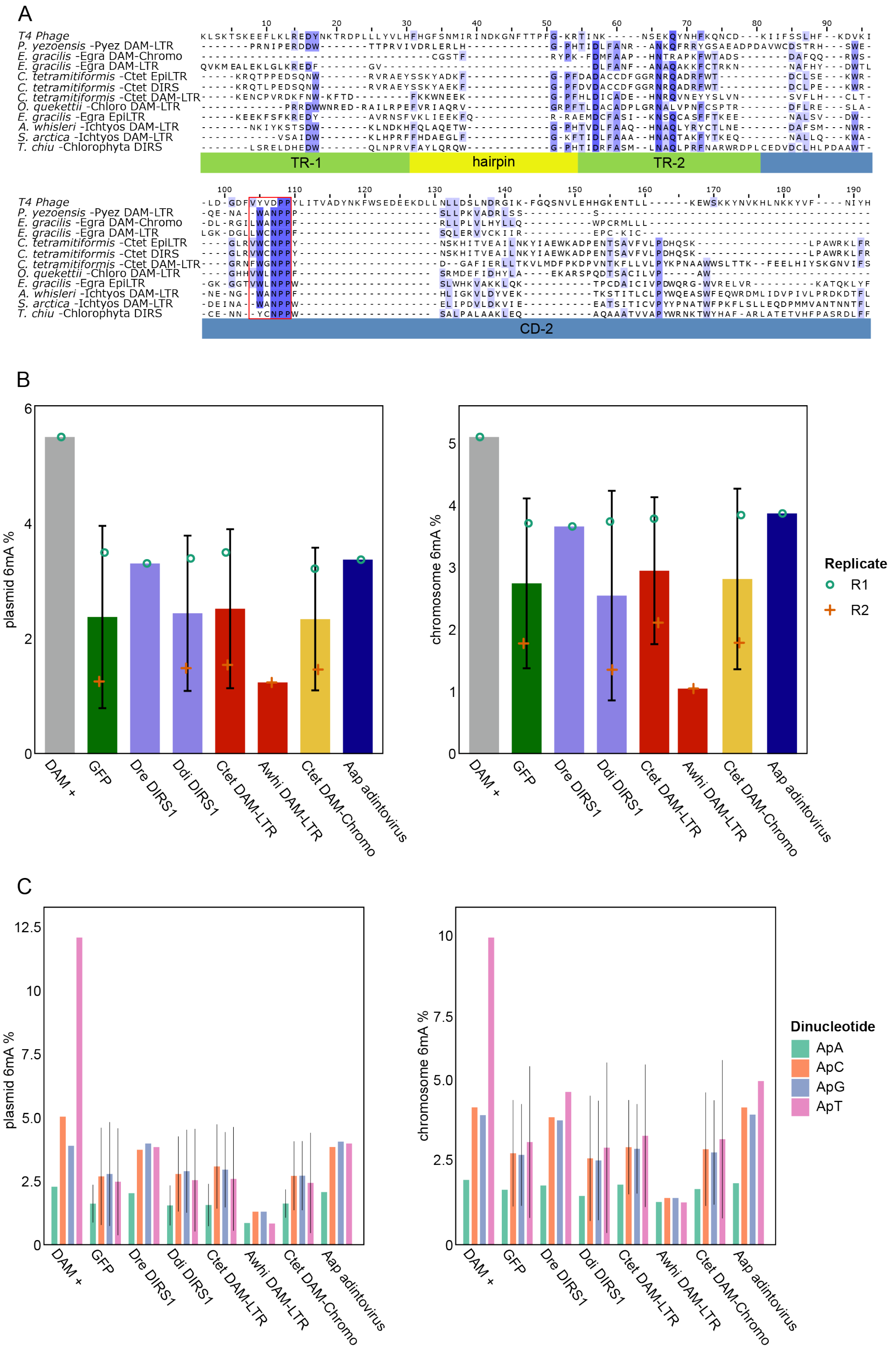


**Supplementary figure 3. DAM conservation and *E. coli* expression.** a) Alignment of representative examples of DAM-TEs and the well-studied T4 phage DAM protein. b) Global 6mA percentage assessed using Oxford Nanopore in plasmid and chromosome of wild type *E. coli* (DAM +) and DAM deficient *E. coli* expressing GFP (as negative control) and a set of representative eukaryotic DAM domains: *Danio rerio* DIRS1 (Dre DIRS1), *Dictyostelium*

*discoideum* DIRS1 (Ddi DIRS1), *Cymbomonas tetramitiformis* DAM-LTR (Ctet DAM-LTR) and DAM-Chromo-LTR (Ctet DAM-Chromo), *Abeoforma whisleri* DAM-LTR (Awhi DAM-LTR), and *Amoebidium appalachense* adintovirus. c) Same but dividing 6mA levels by specific dinucleotide contexts.

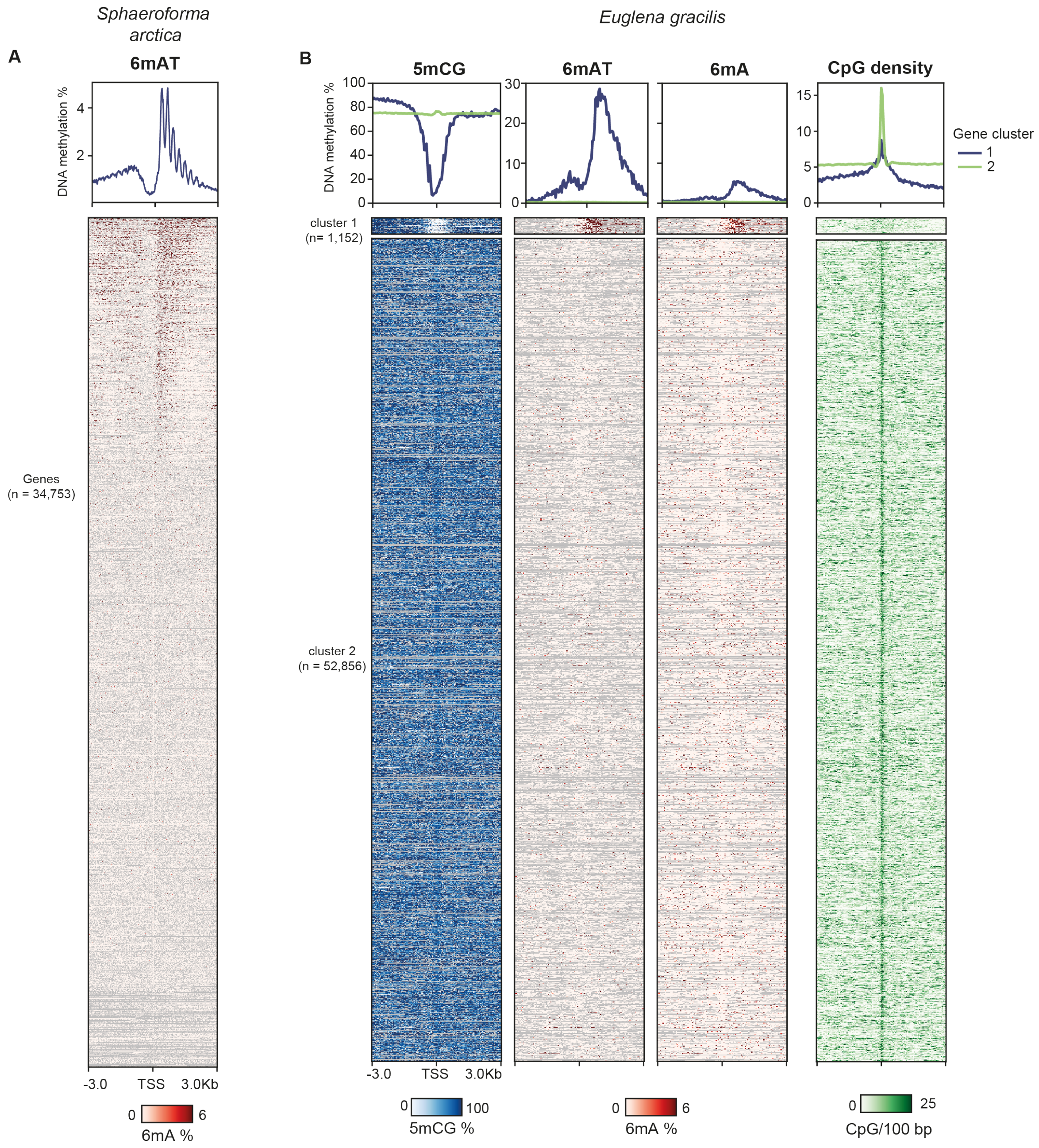


**Supplementary Figure 4. Epigenome profiles for *S. arctica* and *E. gracilis*.** a) Heatmap and average 6mA methylation levels around the Transcriptional Start Site (TSS) of protein coding genes in the ichthyosporean *S. arctica*, highly resembling those of other ichthyosporeans. b) Heatmap and average 5mCG, 6mA (ApTs and all As) and CpG density levels around the TSS of *E. gracilis*. Genes were clustered using k-means = 2 in PlotHeatmap function. CpG density represents the number of CpGs per 100 bp windows.

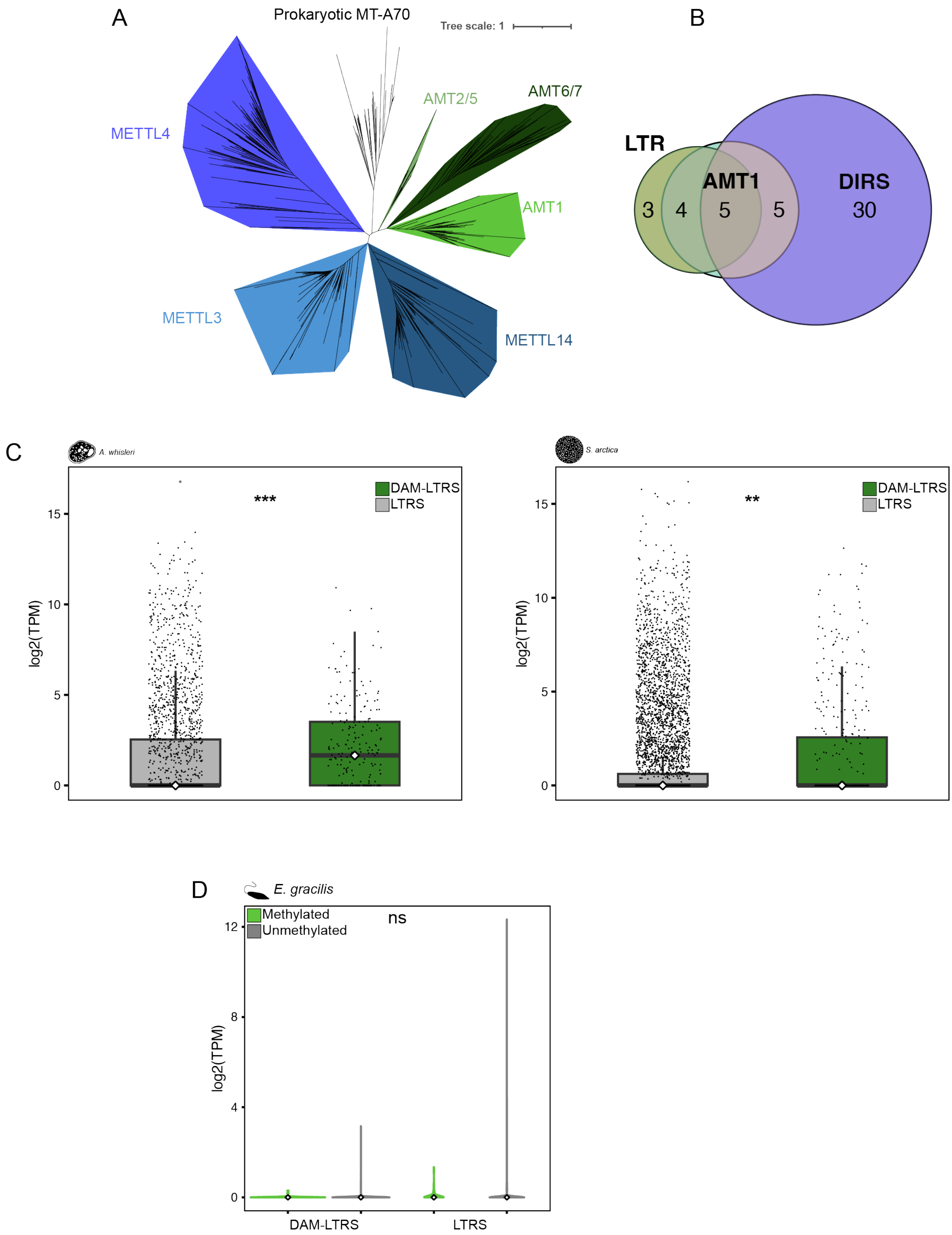


**Supplementary Figure 5. DAM-TE adenine methylation and transcriptional activity.** a) maximum likelihood phylogenetic tree of MT-70 domains used for co-occurrence analysis. b) Venn diagram of species containin DAM-LTRs, DAM-DIRS and AMT1. c) Comparative expression of DAM-LTRs in *A. whisleri* and *S. arctica*. d) Comparative expression of DAM-LTRs and LTRs by methylation level in *E. gracilis*.
